# Chikungunya virus non-structural protein 2 (nsP2) inhibits RIG-I and TLR-mediated immune response

**DOI:** 10.1101/2025.04.30.651505

**Authors:** Shivani Raj, Minnah Irfan, C.T. Ranjith-Kumar

## Abstract

Chikungunya virus (CHIKV) is a single-stranded positive-sense RNA virus that employs various strategies to evade the host’s immune response. The CHIKV non-structural protein 2 (nsP2) is one of the viral-encoded proteins essential for viral replication as well as modulation of the immune response. In this study, we demonstrate that CHIKV nsP2 suppresses both RIG-I-like receptors (RLR) and Toll-like receptors (TLR) signaling pathways. Domain mapping of the nsP2 identified the N-terminal region encompassing the N-terminal domain and the helicase domain (NH) is responsible for the inhibition. Furthermore, site-directed mutagenesis experiments showed that functional helicase is necessary for inhibiting interferon production, but the C-terminal V-Loop region, previously implicated in host transcriptional shutoff is not. Lastly, we demonstrate that nsP2 disrupts two key immune signaling pathways, RLR and TLRs, by interfering with common proteins, TBK1 and IRF3, involved in both signaling pathways. These findings enhance our understanding of CHIKV immune evasion strategies and offer potential targets for the development of antiviral therapeutics.

## Introduction

CHIKV belongs to the *Alphavirus* genus within *Togaviridae* family and causes acute febrile illness known as chikungunya fever in humans. It is an arthropod-borne virus primarily transmitted by *Aedes albopictus* and *Aedes aegypti* mosquitoes (Vairo et al., 2019). CHIKV has a positive-sense, single-stranded RNA genome of 11.8 kb. The CHIKV genome contains two open reading frames (ORFs). ORF1 is processed into four non-structural proteins (nsP1, nsP2, nsP3, and nsP4) essential for virus replication and RNA transcription. While ORF2 is processed into structural proteins (C, E1, E2, E3, and 6K protein) (Silva and Dermody, 2017).

The innate immune response is the first line of defense against viral infection. Activation of the immune response leads to the production of interferons and cytokines, which limit the replication of the virus (Rai et al., 2021). Viral proteins not only play a crucial role in replication but also use multiple strategies to disrupt cellular mechanisms, allowing viruses to evade immune response and establish infection. Detection of viral RNA or pathogen-associated molecular patterns (PAMPs) is mediated by pathogen recognition receptors (PRRs), like TLRs and RLRs (Carty et al., 2021; Okamoto et al., 2017; Wicherska-Pawłowska et al., 2021; Yoneyama et al., 2015). These PRRs recognize viral RNA replication intermediates that trigger downstream signaling via adaptor proteins like mitochondrial antiviral signaling (MAVS) for RLRs, while TLRs use TIR-domain-containing adaptor inducing interferon β (TRIF) and myeloid differentiation primary response gene 88 (MyD88) (Kawai and Akira, 2008; Wicherska-Pawłowska et al., 2021). This triggers a cascade of signaling events that activate TANK-binding kinase 1 (TBK1), which further phosphorylates the interferon regulatory factor 3 (IRF3), ultimately leading to the production of interferons and cytokines (Fitzgerald et al., 2003).

Like many alphaviruses, CHIKV uses both non-structural and structural proteins to evade the immune response (Liu et al., 2022). nsP2 is a multifunctional protein necessary for viral replication and is known to interfere with the immune response (Wang et al., 2025). The N-terminus of CHIKV nsP2 possesses nucleoside triphosphatase, helicase, and RNA-dependent 5’-triphosphatase activity, while the C-terminus harbors a protease domain and methyltransferase-like domain (MTase) (Fros et al., 2013; Karpe et al., 2011). Structurally, the N-terminal region is further classified into the N-terminal domain (NTD), Stalk α-helix, and accessory domain 1B, all of which wrap around the conserved helicase core, ensuring conformational stability (Law et al., 2019). Notably, the first 14 amino acids of the N-terminal region play a crucial role in stabilizing the structure and helicase activity; disruption in this region results in impairment of the function (Das et al., 2014; Law et al., 2019).

The helicase domain is involved in unwinding of viral RNA during replication and transcription (Ranji and Boris-Lawrie, 2010). Mutations within conserved helicase motifs significantly impair its functioning (Grimes and Denison, 2024; Hall and Matson, 1999; Law et al., 2021). CHIKV nsP2 helicase is a member of the helicase superfamily 1 (SF1) and contains seven conserved signature motifs (I, Ia, II, III, IV, V, and VI), which form the enzyme’s catalytic core (Singleton et al., 2007). The Walker A motif is characterized by lysine residues and binds the β- and γ-phosphates of NTP-Mg^2^□, while the Walker B motif contains a conserved aspartate residue that interacts with Mg^2^□ (Das et al., 2014; Karpe et al., 2011). In addition, the C-terminal region of the nsP2 harbors a segment called V-Loop (residues A674, T675, and L676), which has been shown to degrade RNA polymerase II subunit Rpb1, leading to host transcription shutoff (Akhrymuk et al., 2012).

Thus, CHIKV employs various strategies to evade immune response, with nsP2 playing a crucial role in suppressing innate immunity. However, the precise molecular mechanism underlying this immune evasion is not well understood. In this study, we sought to elucidate the role of nsP2’s functional domains in immune inhibition by generating various deletion mutants. Furthermore, we introduced targeted mutations in the conserved Walker motifs to disrupt helicase activity, as well as in the C-terminal V-Loop region, previously implicated in host transcriptional shutoff. Our findings indicate that the N-terminal domain and helicase domain (NH) of the protein are crucial for nsP2-mediated immune suppression and are not dependent on host transcriptional shutoff, highlighting a novel functional aspect of this viral protein. By uncovering the mechanisms through which nsP2 impairs interferon and pro-inflammatory cytokine production, this study enhances our understanding of CHIKV pathogenesis and provides a foundation for the development of targeted antiviral strategies.

## Materal and methods

### Plasmid construct

Plasmids encoding C-terminal FLAG-tagged CHIKV non-structural proteins (nsP1, nsP2, nsP3, nsP4), subdomains of nsP2 (NTD, NH, H, HPM, NHP, HP, P, M), and deletion constructs of NH (Δ14 NH, Δ77 NH, Δ110 NH were generated by subcloning into pUNO (Invivogen) expression vector. Helicase mutants (K192A, D252A, and DE252/253AA) were generated using site-directed mutagenesis and cloned into pUNO-MCS. Similarly, RLE mutant (V-Loop) was also generated. Clones were generated by PCR using CHIKV strain IND-06-Guj as a template. Both the vector pUNO and the PCR products were digested using *AgeI* and *NheI* restriction enzymes. The digested PCR products and vector were ligated, and the ligated products were transformed and expressed in *E*.*coli* Top10 cells. Further clones were subjected to full-length DNA sequencing. Primer sequences are listed in the Supplementary Table 1 (S1).

### Maintenance of Cell lines

HEK293T cells and Huh7 cells were maintained in Dulbecco’s Modified Eagle Medium (DMEM) supplemented with 10 % Fetal Bovine Serum (FBS) and penicillin/streptomycin at 37 °C, 5% CO2. For transfection, cells were seeded at 70-80% confluency.

### Dual-Luciferase Assay

The day prior to transfection, HEK293T cells (40,000 cells/well) were seeded into a CoStar White 96-well plate. Cells were co-transfected with Firefly luciferase (30ng) cloned under the IFN-β promoter (IFNβ-luc) was used as the reporter to measure the IFN-β promoter induction. Renilla luciferase (5ng) cloned under thymidine kinase promoter (phRLTKLuc) was used as an internal control. To induce RIG-I signaling, RIG-I (10ng) was co-transfected with the plasmids of nsPs, which were activated using 3pdsRNA post 24 hours of transfection. For further experiments, CARD domain mutant (R-C) was used instead of RIG-I as its overexpression mimics RIG-I signaling. R-C (10ng) was co-transfected along with nsP2 and its deletion mutant plasmids (25ng). For IPS-1 assay IPS-1 plasmid (10ng), for MyD88 assay-for TRIF assay-myc-TRIF (25ng), MyD88 (25ng), for TBK1 assay-HA-TBK1 (25ng), and for IRF3 assay-HA-IRF3 (25ng) were transfected along with the required plasmids (25ng) as described above. According to the manufacturer’s protocol, HEK293T cells were transfected with varying amounts of plasmids using Lipofectamine 2000. Promega DualGlo Luciferase assay system reagents were used to detect the luciferase activity 48 hours after transfection.

### Cytotoxic Assay

40,000 HEK293T cells were seeded per well into a CoStar White 96-well plate and grown overnight at 37 °C. After 16h, cells were transfected with different amounts of nsP2 (0 to 200ng) plasmid using Lipofectamine 2000 according to the manufacturer’s protocol. After 24h, the media was removed, and Roche WST-1 reagent was used to measure the viability of cells. Absorbance was measured at 450nm and 630 nm using an ELISA absorbance reader.

### RT-PCR

To analyze the expression of downstream IFNβ induced by R-C signaling by Real-time PCR. HEK293T cells were seeded in 12-well plate with 400,000 cells per well, one day before the transfection. The next day, cells were transfected with the R-C in the presence and absence of nsP2 using Lipofectamine 2000 according to the manufacturer’s protocol. R-C (100ng) was co-transfected along with nsP2 and its deletion mutant plasmids (250ng). For IPS-1 assay IPS-1 plasmid (100ng), for MyD88 assay-for TRIF assay-myc-TRIF (250ng), MyD88 (250ng), for TBK1 assay-HA-TBK1 (250ng), and for IRF3 assay-HA-IRF3 (250ng) were transfected along with the required plasmids (250ng) as described above. For dose-dependent assay nsP2 and NH (10 to 250ng) was transfected along with R-C. 48h post-transfection, total RNA was isolated using RNAiso Plus reagent (Takara Bio Companies) according to the manufacturer’s protocol. cDNA synthesis was performed for the total RNA isolated using random hexamers according to the manufacturer’s protocol (PrimeScript 1st strand cDNA synthesis kit, Takara Bio Companies). Quantitative Real-Time PCR (qRT-PCR) was carried out using gene-specific primers to quantify mRNA levels of IFNβ by TB green (Takara Bio Companies). Ct values corresponding to IFNβ mRNA levels measured and normalized with GAPDH Ct values and change in mRNA levels was calculated by ΔΔCt method with respect to vector control.

### Western Blotting

HEK293T cells were transfected with the plasmids of clones generated. Post 48h transfection, transfected cells were washed with the 1ml 1X PBS, lysed with Laemmli buffer, and kept at 95oC for 10 mins. Lysates of transfected cells were loaded into well of 12% polyacrylamide gel. SDS-PAGE separated the proteins and then transferred them to a nitrocellulose membrane and detected them using an anti-FLAG antibody (1:2000). Detection of the antibody-protein complex was done using an ECL western blotting kit according to the manufacturer’s protocol.

### Statistical analysis

All data were analyzed using GraphPad Prism software. Statistical analysis between groups was conducted using a t-test. The differences among samples were considered significant for the values <0.05.

## Results

### nsP2 inhibits the RIG-I signaling

To assess whether any of the CHIKV non-structural protein(s) interferes with the RIG-I signaling, each of the nsPs (nsP1, nsP2, nsP3, and nsP4) were cloned into the mammalian expression vector pUNO-MCS [Fig. 1A], and their expression was confirmed by western blotting using anti-FLAG antibody [Fig. S1A]. The effect of CHIKV nsP(s) on RIG-I signaling was determined by dual luciferase reporter assay as described earlier (Ranjith-Kumar et al., 2009). Briefly, HEK293T cells were co-transfected with each of the nsPs along with the plasmid expressing full-length RIG-I receptor and luciferase reporter plasmids. 24h post-transfection, 5’ triphosphorylated dsRNA of 27bp (3PdsRNA) was used to activate RIG-I signaling (Lu et al., 2010). The reporter plasmids expressing Firefly luciferase under the IFN-β promoter (IFN-β-luc) was used to measure IFN-β promoter activation, and Renilla luciferase under the thymidine kinase promoter (phRLTKLuc) used as an internal control (Ranjith-Kumar et al., 2010). Among all non-structural proteins of CHIKV, nsP2 significantly suppressed RIG-I mediated signaling [Fig. 1B]. To analyze whether nsP2 specifically targets RIG-I receptor or its downstream signaling, we transfected the constitutively active mutant of RIG-I, consisting of only its CARD domains (R-C) instead of full-length RIG-I [Fig. 1C]. Overexpression of R-C was reported to induce an interferon response in the absence of an agonist (Kumar et al., 2023). Hence, R-C was co-transfected along with individual non-structural proteins, and similar to the inhibition of 3PdsRNA-induced RIG-I signaling, nsP2 inhibited R-C induced signaling [Fig. 1C]. This suggests that nsP2 could either be blocking the CARD domains of RIG-I or acting at a step further downstream. Further, a WST-I (Water Soluble Tetrazolium 1) assay was carried out to confirm that the observed inhibition was not due to any cytotoxicity due to overexpression of nsP2 [Fig. 1D]. No cytotoxic effect was observed, further confirming that nsP2’s inhibitory effect on RIG-I signaling was not due to any cell death. Additionally, RT-qPCR analysis was performed to determine the effect of nsP2 at the transcriptional levels of IFN-β, which as expected, revealed a significant reduction in the IFN-β mRNA, confirming the nsP2-mediated inhibition of RIG-I signaling [Fig. 1E]. These data suggest that nsP2 is involved in inhibiting the RIG-I signaling pathway.

**Figure 1.**
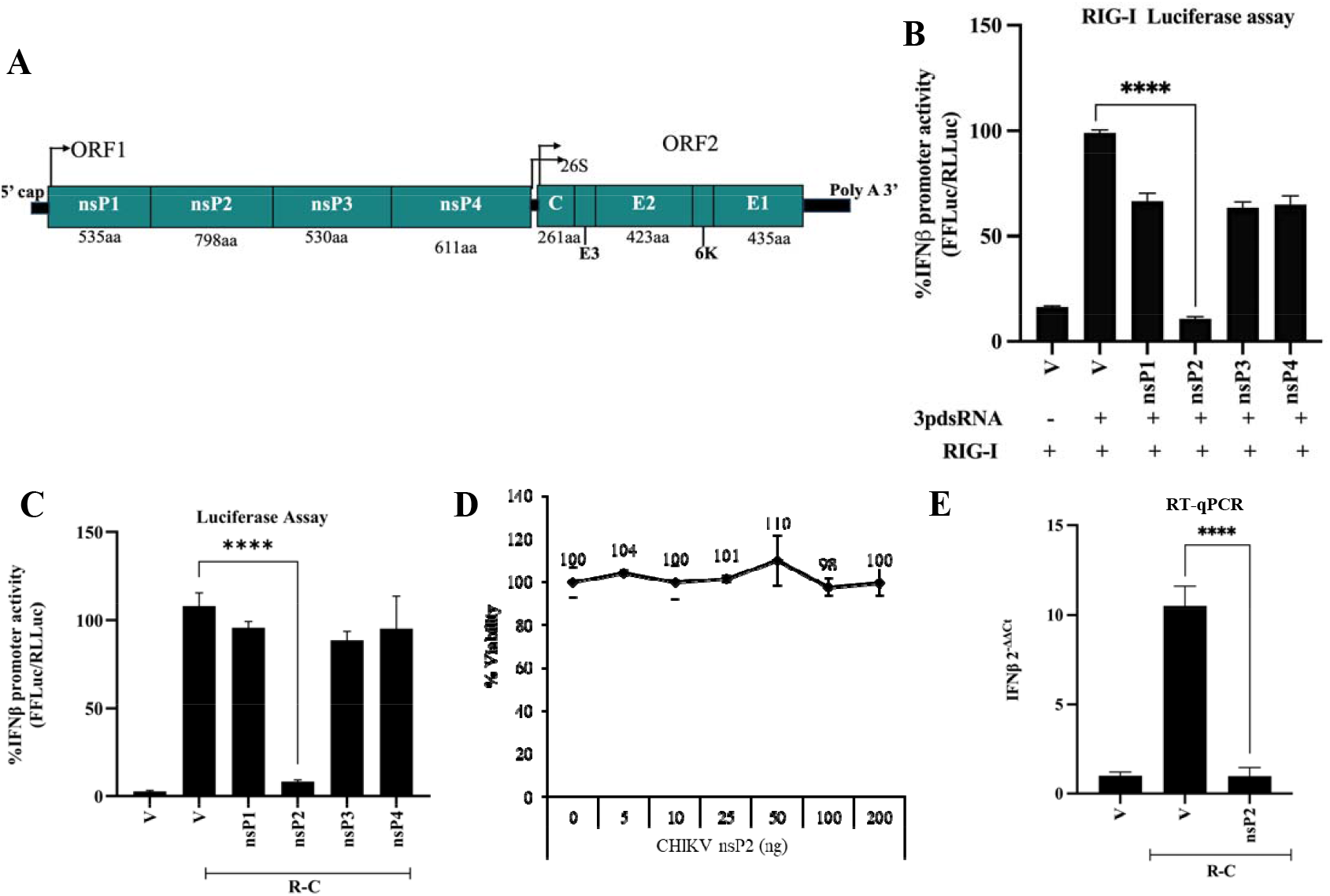
CHIKV non-structural protein 2 (nsP2) inhibits RIG-I signaling. (A) Schematic representation of the CHIKV genome depicting nsPs and SPs. (B) Dual luciferase assay to analyze RIG-I mediated IFN-β promoter activity, plasmids expressing nsP1, nsP2, nsP3, nsP4 and RIG-I co-transfected along with IFN-β firefly and TK *Renilla* luciferase reporters in HEK293T cells, RIG-I was induced with 3p27dsRNA 24h post-transfection. Luciferase activity was measured post-16h induction. (C) Dual luciferase assay to analyze R-C mediated IFN-β promoter activity, plasmids expressing nsP1, nsP2, nsP3, nsP4, and R-C constitutively active CARD domain were co-transfected along with IFN-β firefly and TK *Renilla* luciferase reporters. (D) Cell viability assay-nsP2 plasmid at different amounts from 0 to 200 ng, transfected post 48h of transfection using WST-1 reagent, absorbance measured at 450nm. Values were normalized to control absorbance measured at 630nm. (E) RT-qPCR to analyze IFN-β mRNA levels, HEK293T cells were co-transfected with R-C and nsP2 in a 12-well plate. Post 48h of transfection, RNA was isolated. Values are 2^-ΔΔCt^ ± SD for RT-qPCR and mean percent ± SD for luciferase assay. n=3 for all experiments (**** denotes p value ≤ 0.001).

### VLoop region of the C-terminal domain of nsP2 is not necessary for the inhibition of innate immune response

The C-terminal VLoop region (amino acids 674 to 676) of nsP2 was previously implicated to mediate host transcriptional shutoff by targeting the catalytic subunit of RNAPII, Rpb1, for degradation (Akhrymuk et al., 2018; Meshram et al., 2019). Multiple sequence alignment using Clustal Omega (Madeira et al., 2024) revealed conservation of VLoop (ATL) residues among few alphaviruses [Fig. 2A]. To explore whether transcriptional shutoff by C-terminal VLoop (ATL) contributes directly to the observed nsP2-mediated inhibition of RIG-I signaling, mutation was introduced in the VLoop (A647R, T675L, L767E: referred to as RLE mutant) region of full-length nsP2 based on the findings of Akhrymuk et al., 2019 (Akhrymuk et al., 2019). To assess potential structural change caused by this mutation, we predicted and compared the structure of WT-nsP2 and nsP2 RLE mutant using AlphaFold (Abramson et al., 2024). As shown in Fig. 2B, the RLE mutation disrupted the structure of the VLoop region. To evaluate whether structural and functional alteration of nsP2 impacts RIG-I signaling, the RLE mutant was constructed in pUNO-MCS, and protein expression was confirmed by western analysis [Fig. S1B]. To determine its effect on RIG-I signaling, IFN-β promoter activation and mRNA levels were measured by dual luciferase reporter assay and RT-qPCR, respectively [Fig. 2C & 2D]. Interestingly, RLE mutant showed similar inhibition of RIG-I signaling compared to wild-type nsP2. Despite lacking transcriptional shutoff activity, RLE effectively inhibited RIG-I signaling, indicating that nsP2 employs a distinct mechanism to interfere with the immune response that functions independently of host transcriptional shutoff. To further understand the mechanism, we aimed to determine the specific domains of nsP2 that are involved in mediating the inhibition.

**Figure 2.**
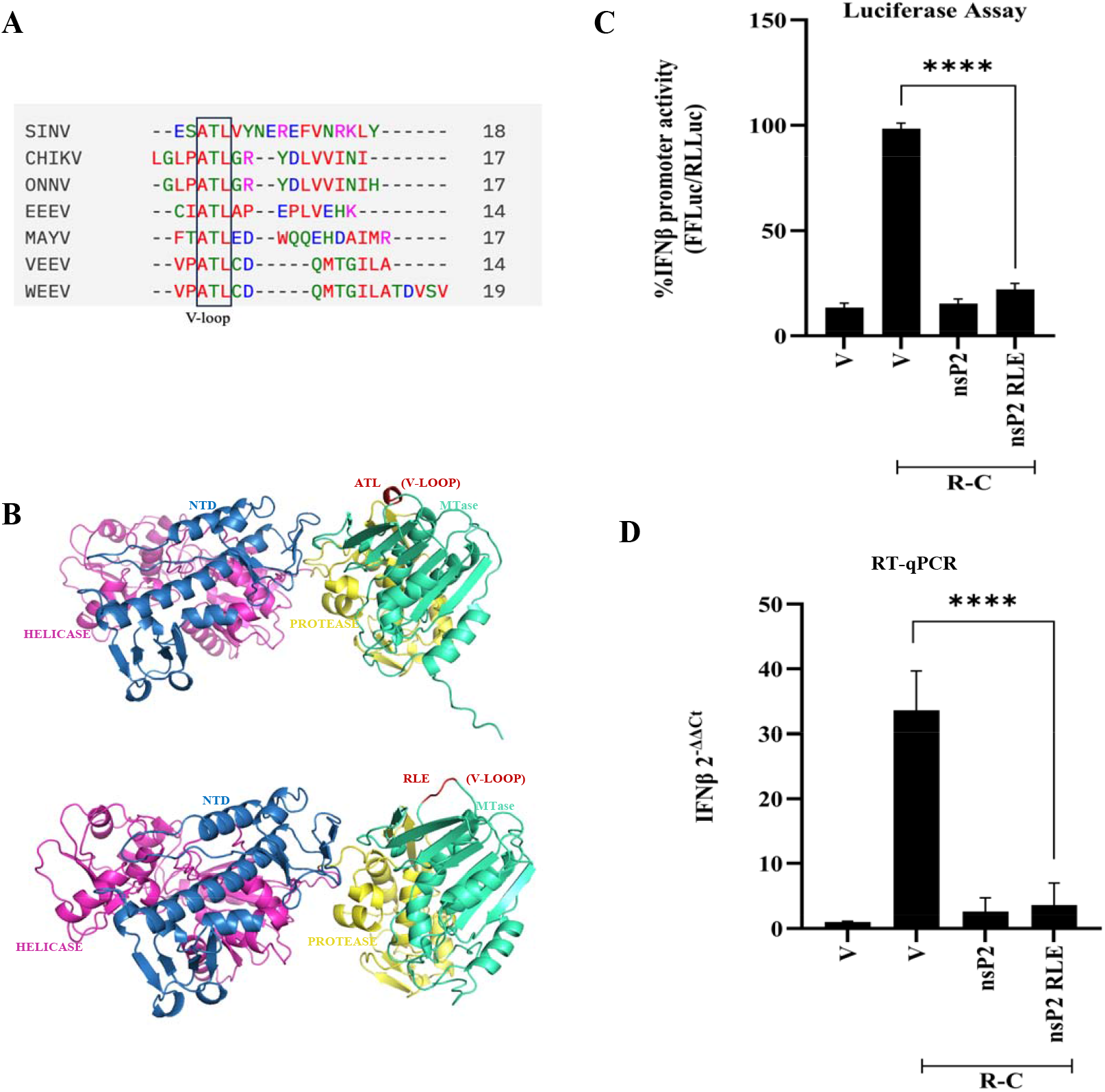
nsP2 RLE mutant does inhibit RIG-I signaling (A) The sequence of CHIKV nsP2 C-terminal domain (605 aa - 798 aa) aligned with the C-terminal of Sindbis virus (SINV), O’nyong’nyong virus (ONNV), Eastern Equine Encephalitis virus (EEEV), Venezuelan equine encephalitis virus (VEEV), Western equine encephalitis virus (WEEV), and Mayaro virus (MAYV). The highlighted conserved residues in all the sequences indicate a V-loop (ATL) region. (B) The predicted 3D structure of nsP2 using the Alphafold server shows the NTD (blue), Helicase (magenta), protease (yellow), and MTase domain (cyan). The ATL (V-Loop) of nsP2 is highlighted in red and mutated into RLE. (C) Dual luciferase assay to analyze R-C mediated IFN-β promoter activity, nsP2, nsP2 RLE, and R-C along with IFN-β firefly and TK *Renilla* luciferase reporters co-transfected in HEK293T cells, and Luciferase activity was measured post 48h transfection. (D) RT-qPCR to analyze IFNβ mRNA levels, HEK293T cells co-transfected with R-C, nsP2, and nsP2 RLE in a 12-well plate. Post 48h of transfection, RNA was isolated. Values are 2^-ΔΔCt^ ± SD for RT-qPCR and mean percent ± SD for luciferase assay. n=3 for all experiments (**** denotes p value ≤ 0.001).

### NH region of CHIKV nsP2 is involved in the inhibition of RIG-I signaling

In order to identify the specific region of nsP2 responsible for RIG-I inhibition, following deletion mutants of nsP2-NTD (1-167 aa), H (167-470 aa), NH (1-470 aa), HP (167-605 aa), NHP (1-605 aa), HPM (167-798 aa), P (470-605 aa), PM (470-798aa), and M (605-798 aa) were constructed [Fig. 3A]. Protein expression was confirmed by western blotting using an anti-FLAG antibody [Fig. S1C]. For analyzing the effect of subdomains of nsP2 on RIG-I signaling, IFN-β promoter activity was measured through dual luciferase reporter assay [Fig. 3B]. The N-terminal domain, along with the helicase domain (NH) region, showed similar inhibition of RIG-I signaling as that of a full-length nsP2. NTD and H domains by themselves showed much reduced inhibition, suggesting that the presence of the two domains together is important for efficient inhibition. Interestingly, the other domains, including NHP, showed significantly less inhibition. It is possible that in the NHP construct, the P domain is interfering with the inhibitory activity of the NH domain, which evidently may not be the case in full-length nsP2. These data suggest that NH might be the primary region of nsP2 necessary for inhibition. A RT-qPCR was performed to measure IFN-β mRNA levels to verify the observed inhibition by nsP2 and NH. As expected, the NH domain showed similar inhibition as full-length nsP2, further confirming that the NH domain within nsP2 is sufficient for inhibiting RIG-I signaling [Fig. 3C]. To determine whether nsP2 also interferes with NF-κB signaling, we measured NF-κB mRNA levels by RT-qPCR. Notably, both nsP2 and NH suppressed NF-κB signaling, suggesting nsP2 broadly targets innate immune signaling downstream of RIG-I [Fig. 3D]. Lastly, to assess the potency of observed inhibition, HEK293T cells were co-transfected with increasing amounts of nsP2 and NH (10ng to 250ng) plasmids along with R-C [Fig. 3E]. At lower amounts, nsP2 was more efficient in inhibiting RIG-I signaling compared to NH, suggesting that while NH is an essential region required for inhibition of the immune response, additional domains may enhance the efficiency of inhibition by providing structural stability or facilitating interaction.

**Figure 3.**
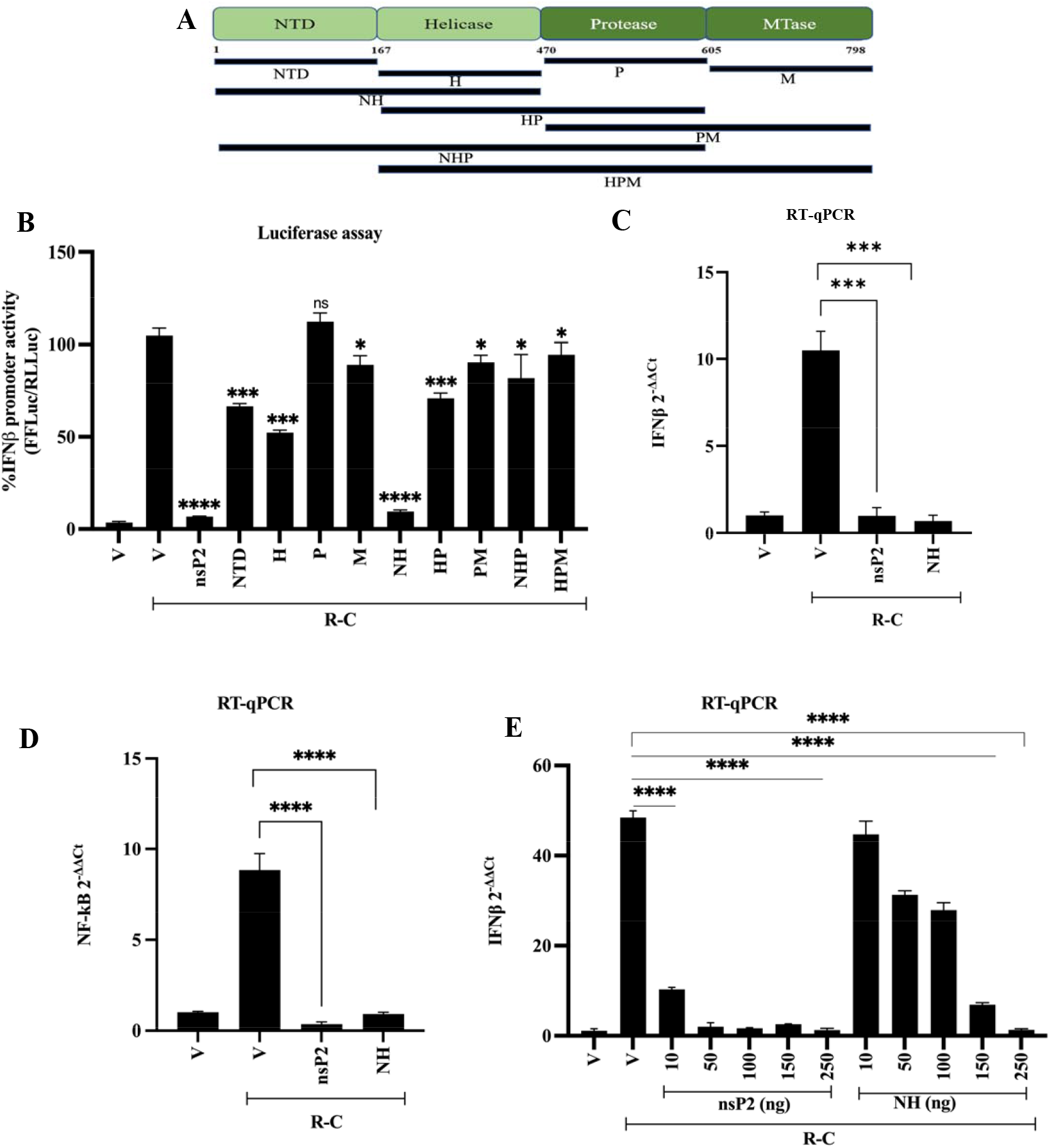
NH (1-470 aa) of nsP2 inhibits RIG-I signaling (A) Schematic representation of CHIKV nsP2 subdomains, (B) Dual luciferase assay to analyze R-C mediated IFN-β promoter activity, nsP2, subdomains of nsP2, and R-C along with IFN-β firefly and TK *Renilla* luciferase reporters co-transfected in HEK293T cells, and Luciferase activity was measured post 48h transfection. (C) RT-qPCR to analyze IFN-β mRNA levels, HEK293T cells co-transfected with R-C, nsP2, and NH. Post 48h of transfection, RNA was isolated. (D) Quantitative RT-PCR to analyze NF-κB mRNA levels, HEK293T cells co-transfected with R-C, nsP2, and NH. Post 48h of transfection, RNA was isolated. Ct values corresponding to levels measured and normalized with GAPDH Ct values and change in mRNA levels was calculated by ^ΔΔCt^ method with respect to IFN-β mRNA vector control. [E] For amount-dependent assay, HEK293T cells were transfected with increasing amounts of nsP2 and NH plasmids in a 12-well plate. Post 48h of transfection, RNA was isolated. Values are 2^-ΔΔCt^ ± SD for RT-qPCR and mean percent ± SD for luciferase assay. n=3 for all experiments (**** denotes p value ≤ 0.001 * denotes p value ≤ 0.05).

### Minimal region of NH involved in the inhibition of RIG-I signaling

The N-terminal region is classified into subregions NTD (77aa), stalk (77aa-110aa), and 1B (110aa-175aa), which are the accessory domains of nsP2 necessary for helicase activity (Law et al., 2019). To identify the minimal region of nsP2 necessary for the observed inhibition, further mutants of NH were generated wherein N-terminal 14aa (Δ14 NH), 77aa (Δ77 NH), and 110aa (Δ110 NH) were deleted [Fig. 4A]. The deletion mutants were cloned in pUNO-MCS mammalian expression vector, and their expression was confirmed using western analysis [Fig. S1D]. The impact of these deletion mutants on RIG-I signaling was assessed by dual luciferase reporter assay and was further validated by RT-qPCR [Fig. 4B & 4C]. The N-terminal deletion mutants showed no significant inhibition compared to NH in both dual luciferase and RT-qPCR assays. Our findings demonstrate that the entire N-terminal domain and Helicase domain (NH) might be important for the observed inhibition of RIG-I signaling. Interestingly, it was reported earlier that the N-terminal 14 amino acids of nsP2 are important for the stability and activity of helicase [14].

**Figure 4.**
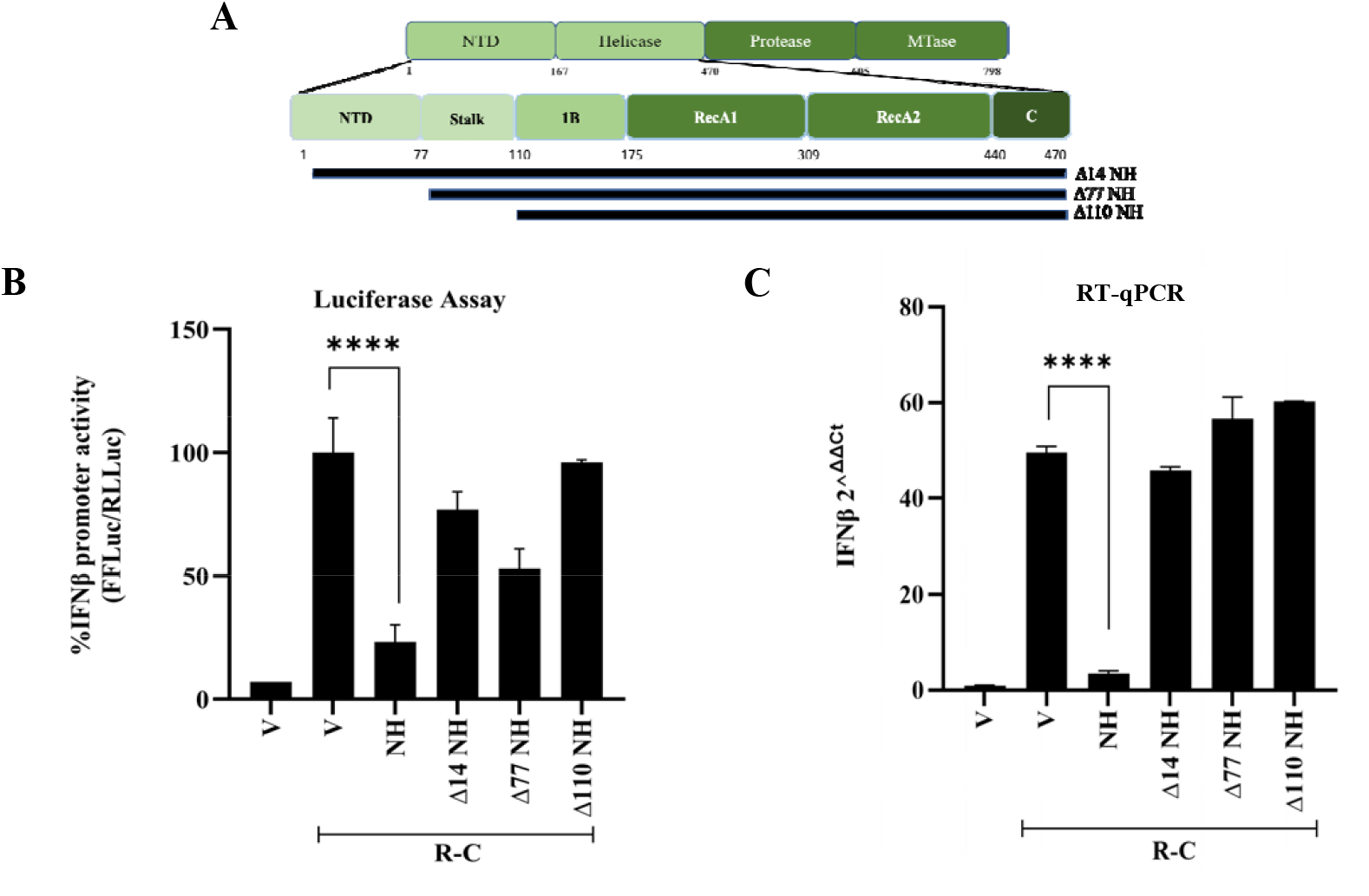
nsP2-NH is the minimal region that inhibits RIG-I signaling (A) Schematic representation of CHIKV nsP2-NH subdomains, (B) Dual luciferase assay to analyze R-C mediated IFN-β promoter activity, NH, Δ14 NH, Δ77 NH, and Δ110 NH, and R-C along with IFN-β firefly and TK *Renilla* luciferase reporters co-transfected in HEK293T cells, and Luciferase activity was measured post 48h transfection. (C) RT-qPCR to analyze IFN-β mRNA levels, HEK293T cells co-transfected with R-C, NH, Δ14 NH, Δ77 NH, and Δ110 NH in 12-well plate. Post 48h of transfection, RNA was isolated. Values are 2^-ΔΔCt^ ± SD for RT-qPCR and mean percent ± SD for luciferase assay. n=3 for all experiments (**** denotes p value ≤ 0.001 * denotes p value ≤ 0.05).

### nsP2 requires helicase activity to inhibit RIG-I signaling

Based on the reported importance of the N-terminal region of nsP2 in stability and helicase activity (Law et al., 2019), we next investigated whether the helicase activity of nsP2 contributes to its ability to inhibit RIG-I signaling. Firstly, we performed comparative sequence analysis of the helicase region among various alphaviruses, which as expected revealed that both Walker A (G/AXXXXGKT/S) and Walker B (DEXX) motifs are highly conserved [Fig. 5A]. We further compared the helicase domain of other RNA viruses, which revealed the Walker A motif to be conserved [Fig. 5B]. These conserved domains were subjected to mutational analysis, wherein Walker A (K192A) and Walker B (D252A, DE252/253AA) motif mutants critical for the helicase function were generated, and their expression was confirmed by western analysis [Fig. S1E]. To assess the role of Walker motif mutants of nsP2 on RIG-I signaling, a dual luciferase reporter assay was performed to measure IFN-β promoter activity [Fig. 5C]. Compared to the wild-type (WT) nsP2, Walker motif mutants showed no inhibition of the RIG-I signaling, suggesting that the helicase activity of nsP2 is important for inhibition. This was further confirmed by analyzing IFN-β mRNA levels through RT-qPCR [Fig. 5D].

**Figure 5:**
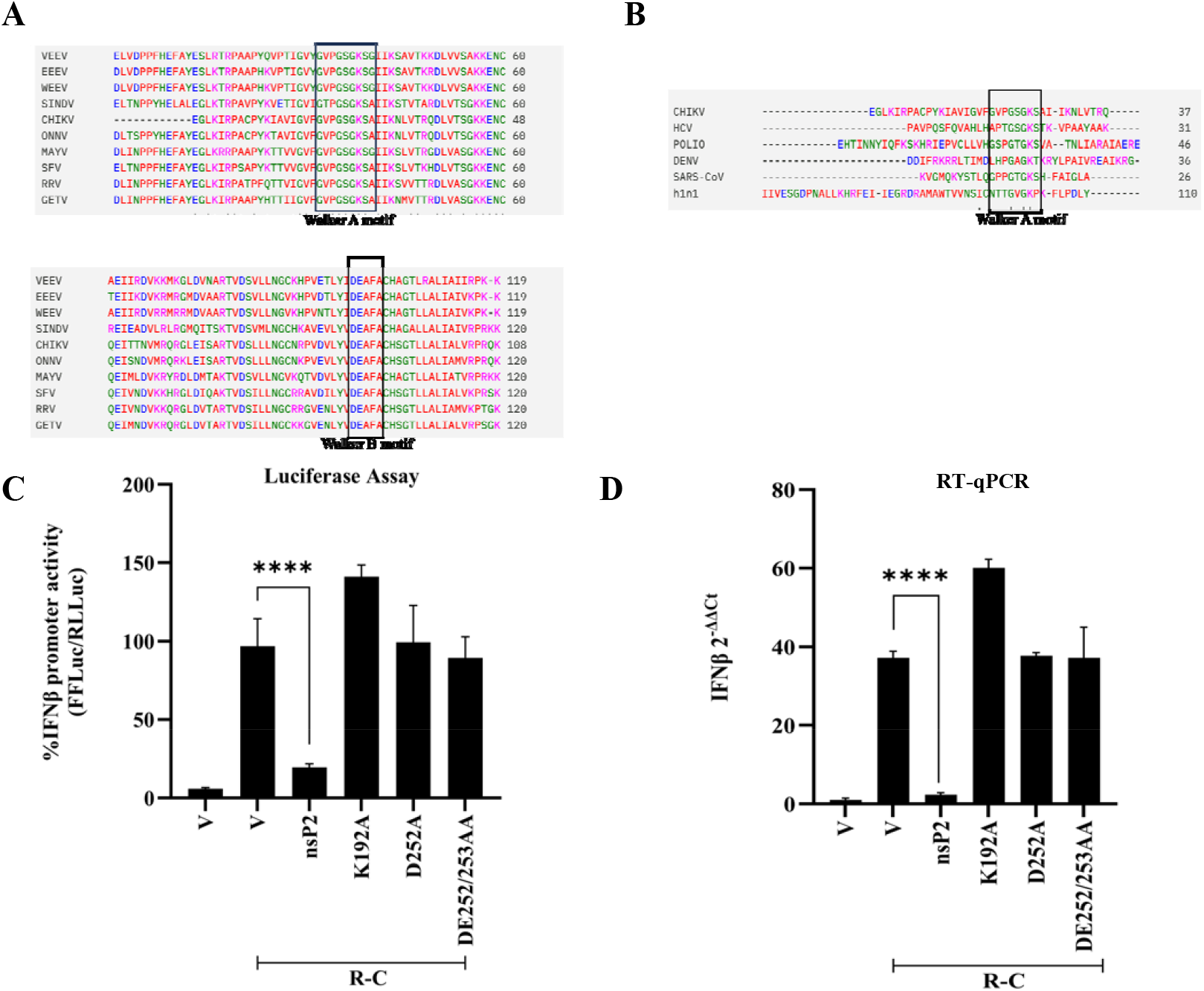
nsP2 helicase mutant does not inhibit RIG-I signaling (A) Using Clustal Omega comparison of sequence encompassing aa residue (167aa-470aa) of CHIKV nsP2 with other alphaviruses-Sindbis virus (SINV), O’nyong’nyong virus (ONNV), Eastern Equine Encephalitis virus (EEEV), Venezuelan equine encephalitis virus (VEEV), Western equine encephalitis virus (WEEV), and Mayaro virus (MAYV). The conserved walker motif regions are highlighted with a box. (B) The comparison of the nsP2 helicase domain with other viruses, Polio Virus, Hepatitis C Virus (HCV), Dengue virus (DENV), SARS-CoV-2, and H1N1 flu (swine flu), shows the conserved region of the Walker A motif. The highlighted helicase Walker motif A (186 aa-193 aa) and Walker motif B (252 aa-255 aa) are then mutated for analysis (C) Dual luciferase assay to analyze R-C mediated IFN-β promoter activity, nsP2, K192A, D252A, and DE252/253AA, and R-C along with IFN-β firefly and TK *Renilla* luciferase reporters co-transfected in HEK293T cells, and Luciferase activity was measured post 48h transfection. (D) RT-qPCR to analyze IFN-β mRNA levels, HEK293T cells co-transfected with R-C, nsP2, K192A, D252A, and DE252/253AA in a 12-well plate. Post 48h of transfection, RNA was isolated. Values are 2^-ΔΔCt^ ± SD for RT-qPCR and mean percent ± SD for luciferase assay. n=3 for all experiments (**** denotes p value ≤ 0.001).

### nsP2 and NH inhibit IPS-I, TRIF, and MyD88-induced IFN-β production

Next, we investigated whether nsP2 and NH target the RIG-I receptor directly to inhibit signaling. For this, we first assessed nsP2’s impact on the adaptor protein of RIG-I signaling, IPS-I (also known as MAVS, Cardiff, VISA) (Potter et al., 2008). Overexpression of IPS-I mimics the activation of RIG-I signaling (Hingane et al., 2020; Kawai et al., 2005). Upon co-expression, both nsP2 and NH suppressed IPS-1 induced signaling significantly in both the dual luciferase reporter assay and RT-qPCR [Fig. 6A and 6D]. In contrast, helicase mutant nsP2-K192A failed to inhibit signaling, further supporting the requirement of functional helicase in nsP2 to inhibit signaling. This also suggests that nsP2 and NH might interfere downstream of RIG-I, rather than interacting directly with the receptor.

**Figure 6:**
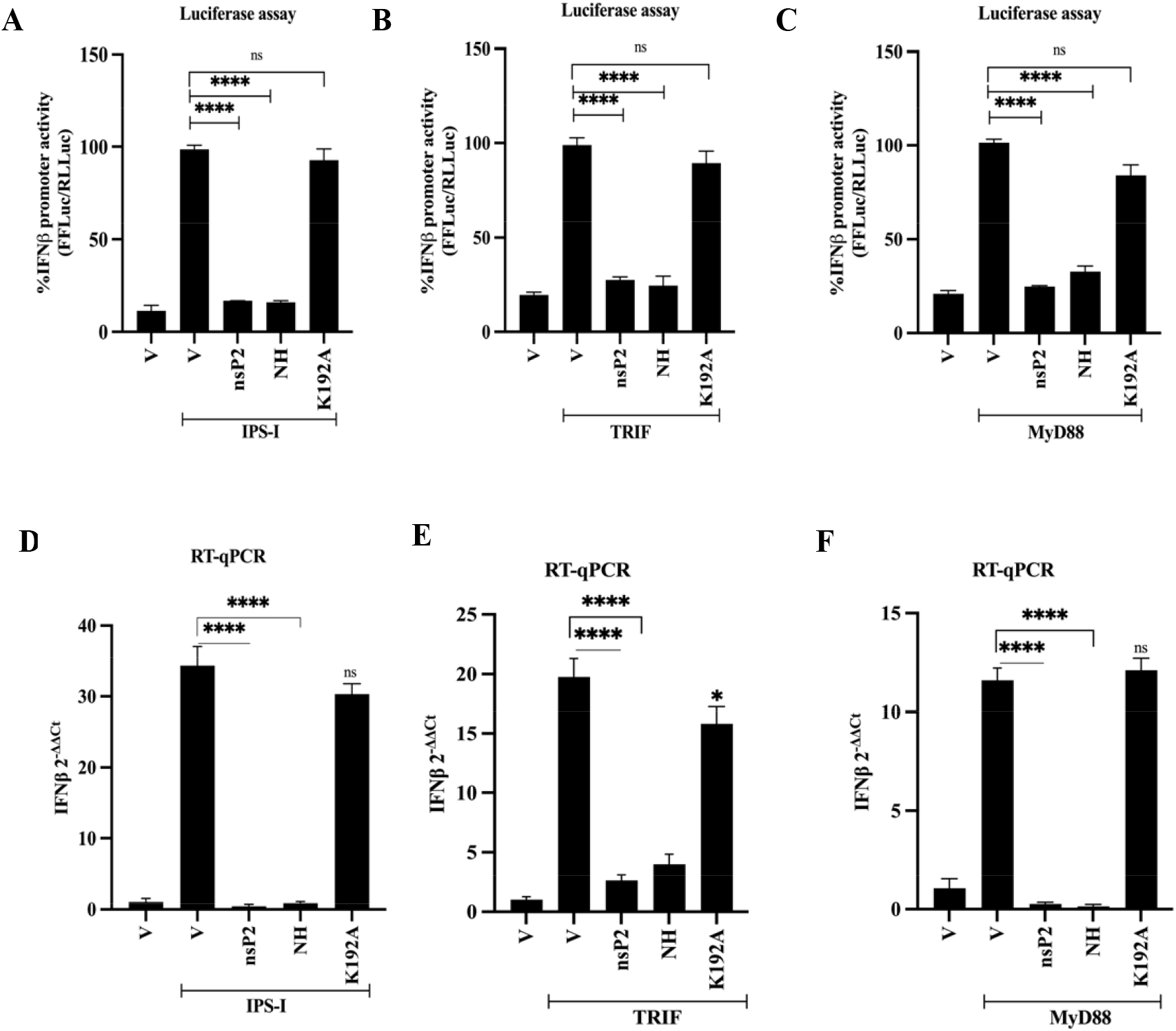
nsP2 inhibits adaptor proteins of RIG-I and TLR signaling. Dual luciferase assay to analyze IPS-I (A), TRIF (B), and MyD88 (C) mediated IFN-β promoter activity, nsP2, NH, helicase mutant K192A, IPS-I, TRIF, MyD88, along with IFN-β firefly and TK *Renilla* luciferase reporters co-transfected in HEK293T cells, and Luciferase activity was measured post 48h transfection. RT-qPCR to analyze IFN-β mRNA levels in IPS-I (D), TRIF (E), and MyD88 (F), HEK293T cells co-transfected with nsP2, NH, K192A, IPS-I, TRIF, and MyD88, in a 12-well plate. Post 48h of transfection, RNA was isolated. Values are mean percent ± SD for the luciferase assay. n=3 for all experiments (**** denotes p value ≤ 0.001). Values are 2^-ΔΔCt^ ± SD for RT qPCR and mean percent ± SD for luciferase assay. n=3 for all experiments (**** denotes p value ≤ 0.001)

To further elucidate whether the observed inhibition is specific to RIG-I signaling, we examined the impact of nsP2 and NH on the adaptor proteins of TLRs, namely TRIF and MyD88. TRIF is the adaptor for TLR3, and MyD88 is the adaptor for all other TLRs (Kawai and Akira, 2008). However, TLR4 uses both TRIF and MyD88 for signaling. Similar to IPS-1, overexpression of these adaptor proteins will mimic TLR signaling (Hingane et al., 2020). nsP2 and NH showed inhibition of the signaling induced by both TRIF and MyD88 in dual luciferase reporter assay [Fig. 6B and 6C] and RT-qPCR [Fig. 6E and 6F], whereas K192A mutant did not exhibit inhibition. This indicates that nsP2 and NH are likely to target downstream common proteins involved in both the TLR and RLR signaling pathways.

### Effect of nsP2 and NH on IRF3 and TBK1

Having shown that nsP2 inhibits both RLR and TLR pathways, we next looked at the possible common proteins involved in these pathways. For this, we investigated the effect of nsP2 on TBK1 and IRF3, which are known to be involved in both RLR and TLR signaling pathways (Fitzgerald et al., 2003). To analyze whether nsP2 and NH target these proteins, IFN-β promoter activation was measured through dual luciferase reporter assay by overexpressing TBK1 and IRF3 in the presence and absence of nsP2 and NH, along with nsP2 helicase mutant K192A [Fig. 7A and 7B]. nsP2 and NH showed significant suppression of TBK1 and IRF3-induced signaling, while K192A failed to show any effect. To further confirm this inhibition at the transcript level, IFN-β mRNA levels were measured by RT-qPCR, nsP2 and NH showed similar inhibition while K192A did not inhibit, suggesting the importance of helicase in the inhibition of IRF3 and TBK1 induced signaling by nsP2 [Fig. 7C and 7D]. This suggests that nsP2 and NH are likely to target common proteins involved in both signaling pathways.

**Figure 7:**
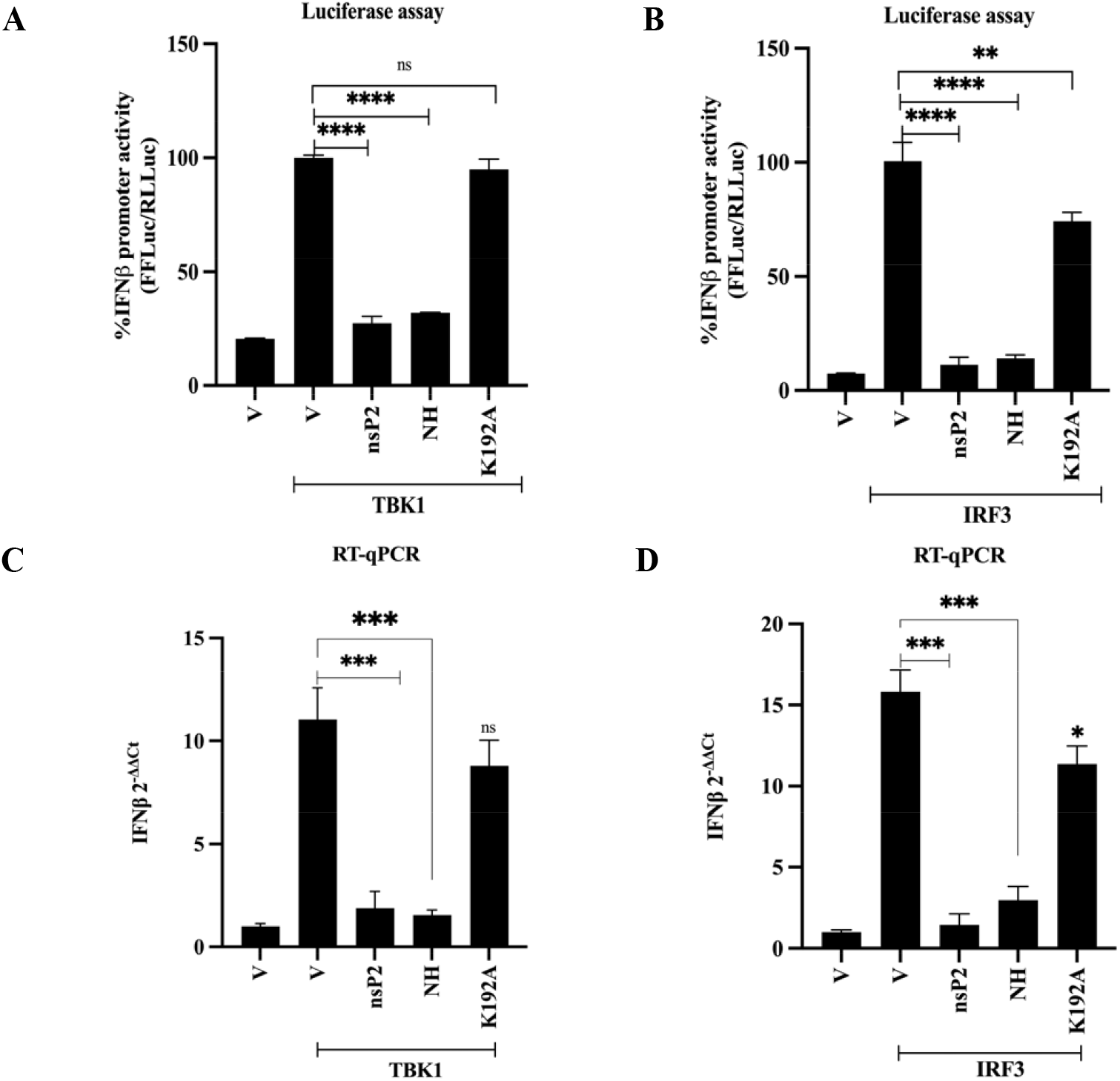
nsP2 and NH inhibit TBK1 and IRF3-induced signaling. (A), (B) Dual luciferase assay to analyze TBK1, IRF3-mediated IFN-β promoter activity, nsP2, NH, helicase mutant K192A, TBK1, IRF3, along with IFN-β firefly and TK *Renilla* luciferase reporters co-transfected. (C), (D) RT-qPCR to analyze IFN-β mRNA levels, HEK293T cells co-transfected with TBK1, IRF3, nsP2, NH, and K192A as negative control in a 12-well plate. Post 48h of transfection, RNA was isolated. Values are 2^-ΔΔCt^ ± SD for RT-qPCR and mean percent ± SD for luciferase assay. n=3 for all experiments (**** denotes p value ≤0.001).

Taken together, our findings demonstrate that the N-terminal domain and the helicase domain (NH) of the nsP2 are sufficient to inhibit both RLR and TLR signaling by targeting common downstream components shared among both signaling pathways, and this inhibition requires a functional helicase domain. Furthermore, our findings imply that nsP2-mediated inhibition occurs independently of host transcriptional shutoff.

## Discussion

The production of IFNs along with pro inflammatory cytokines is a crucial component of the innate immune response against viral infection (Koyama et al., 2008). The PRRs detect PAMPs, such as viral replication intermediates, triggering the production of type I and type III IFNs, along with various cytokines and chemokines (Carty et al., 2021). Among the diverse PRRs, RLRs and TLRs play critical roles in recognizing the viral replication intermediates, including dsRNA, ssRNA, and short 5’ triphosphate RNAs (Lei and Hilgenfeld, 2017; Rai et al., 2021). RNA viruses have evolved various strategies to evade immune detection and modulate signaling pathways, often employing both structural and non-structural proteins to disrupt host immune response and ensure viral survival (Beachboard and Horner, 2016). Understanding these viral mechanisms is crucial for developing antiviral strategies against viral infection.

Previous studies have established that nsP2 of alphaviruses, including CHIKV, not only plays a role in viral replication but also modulates host cellular pathways by interfering with components of immune signaling to evade the host immune response (Akhrymuk et al., 2018; Bae et al., 2019; Meshram et al., 2019). Here we demonstrate that CHIKV nsP2 potently inhibits immune response by suppressing signaling by key adaptor proteins, such as IPS-I (MAVS) for RLR and MyD88 and TRIF for TLRs, suggesting that nsP2 targets common signaling intermediates in these pathways [Fig. 6]. We hypothesized that nsP2 may interfere with common downstream proteins such as TBK1 and IRF3, which are transcriptional regulators essential for the production of interferons and cytokines. Notably, previous findings demonstrated that the ORF2 protein of Hepatitis E Virus (HEV) impedes IRF3 nuclear translocation leading to suppression of IFN production [33]. Our studies revealed that CHIKV nsP2 also suppresses TBK1 and IRF3-mediated signaling [Fig. 7], implying that nsP2 might target these common downstream proteins involved in signaling by either directly targeting them or by interfering with their activation, via signaling complex assembly, phosphorylation events, or nuclear translocation.

Domain mapping of nsP2 further revealed that the N-terminal domain along with helicase domain (NH) of nsP2 is sufficient to mediate the inhibition of RLR and TLR signaling [Fig. 3C & D]. However, NH showed much lower inhibition compared to full-length nsP2 at lower amounts of plasmid [Fig. 3E]. This indicates that full-length nsP2 may exhibit higher affinity or interaction with host target protein(s) involved in these pathways, thereby contributing more effectively to the inhibition of the immune response. This is further substantiated by the observation that domains other than the NH of nsP2 also showed inhibition, though to a much lesser extent [Fig. 3B]. It is also worth noting that NTD and Helicase domains by themselves were unable to show significant inhibition in comparison to NH (harboring both the domains). However, NHP domain was not as potent as NH indicating that the P domain in NHP may be interfering with the ability of NH to inhibit signaling. This also implies a potential structural or organizational difference between NHP and full-length nsP2. Furthermore, deletion analysis of the N-terminal end of nsP2 revealed that the first 14 amino acids are crucial for suppressing RIG-I signaling [Fig. 4]. Previously, this region was reported to be important for NTPase and helicase activities and to stabilize the structure of nsP2 (Law et al., 2019). The need for functional helicase was further substantiated with Walker motif mutants. These findings reveal the importance of helicase activity in the inhibition of RLR and TLR signaling [Figs. 5-7].

Viral helicases are crucial for the unwinding of RNA during replication, but they have also been shown to interfere with the innate immune response (Frick and Lam, 2006; Kwong et al., 2005). In several viruses, helicase domains contribute to immune evasion, like the NS3 helicase of the Hepatitis C virus (HCV), which directly interacts with TBK1, preventing activation of IRF3 and leading to suppression of interferon production (Otsuka et al., 2005). Similarly, Riedl et al., 2019 (Riedl et al., 2019)demonstrated that the helicase domains of NS3 from Zika Virus (ZIKV), Dengue Virus (DENV), and West Nile Virus (WNV) disrupt the interaction between scaffold protein 14-3-3 and RIG-I during antiviral response, resulting in attenuated host immune response (Gack and Diamond, 2016; Riedl et al., 2019). Likewise, the nsp13 helicase of SARS-CoV-2 blocks the interaction of TBK1 and IRF3 to disrupt the signaling (Feng et al., 2023; Fung et al., 2022). Mutational analysis of SARS-CoV-2 nsp13 helicase revealed that a functional helicase is critical for nucleic acid binding. Disruption of this impairs STAT1 phosphorylation, thereby suppressing the immune response (Fung et al., 2022). In addition, NS3 helicase of WNV interferes with the innate immune response by inhibiting STAT1 phosphorylation (Setoh et al., 2017). These findings highlight the role of helicase in immune evasion across viruses. Similarly, our mutational analysis in conserved helicase walker motifs (Walker A and Walker B) of the CHIKV, necessary for ATP hydrolysis and viral RNA replication, prevented nsP2 from inhibiting RIG-I and TLR signaling. These further highlight that helicase activity is essential for the CHIKV nsP2-mediated interference with the immune signaling.

CHIKV nsP2 C-terminal has previously been associated with antagonizing the JAK-STAT signaling pathway and also in promoting the degradation of RNA polymerase II subunit, RbP1, leading to the shutoff of host transcription (Akhrymuk et al., 2012; Bouraï et al., 2012; Fros et al., 2010). Meshram et al., 2019 (Meshram et al., 2019) reported that substitution mutations in the V-Loop, wherein amino acids A, T, and L were substituted with R, L, and E, failed to induce transcriptional shutoff and also did not affect viral replication (Akhrymuk et al., 2019). However, our data demonstrated that the inhibition by nsP2 is not dependent on host transcriptional shutoff as RLE mutant failed to inhibit the RIG-I signaling [Fig. 2]. Furthermore, NH region alone is sufficient to inhibit the RIG-I and TLR signaling, suggesting that the transcriptional shutoff may not be necessary for inhibition of interferon response. This further validates the role of the NH encompassing the active helicase domain of nsP2, which interferes with RLR and TLR signaling. In conclusion, our study shows that CHIKV nsP2 is a major antagonist of the interferon production induced by RLR and TLR signaling pathways. nsP2 likely targets common points downstream of the TLR and RLR signaling to suppress interferon production, although further studies are required to identify the target protein. The requirement for functional helicase and the pivotal role of the NH region in this inhibition provide new insights into the molecular mechanisms. These insights into CHIKV’s evasion of host defense could potentially provide targets for developing antiviral strategies and therapeutics against CHIKV infection.

## Conclusion

Our research provides critical insights into the mechanisms by which CHIKV evades host immune response, specifically through the actions of its non-structural protein 2 (nsP2). While viral proteins are known to suppress innate immune responses to facilitate replication, the precise role of how CHIKV nsP2 interferes remains unclear. Here, we demonstrate that nsP2 disrupts two key immune signaling pathways, RLR and TLRs, by interfering with common proteins involved in both signaling pathways. Through domain mapping and site-directed mutagenesis, we also identified that the N-terminal helicase domain is essential for inhibition. These findings enhance our understanding of CHIKV immune evasion strategies and offer potential targets for the development of antiviral therapeutics.

## Acknowledgments

We thank Professor Sudhanshu Vrati, Regional Center for Biotechnology (RCB), and Dr. Milan Surjit from Translational Health Science and Technology Institute (THSTI) for their valuable help. The authors declare no conflict of interest.

## Funding

This work was funded by the Faculty Research Grant Scheme (FRGS) of Guru Gobind Singh Indraprastha (2022-2025) and was partly funded by the Council of Scientific and Industrial Research (CSIR) (37/1724/19/EMR-II). The Indian Council of Medical Research (ICMR) funded the fellowship of Shivani Raj (Senior Research Fellowship) (VIR/Fellowship/3/2022-ECD-I IRIS No. 2021-14954).

## Notes

### Competing Interest Statement

The authors have declared no competing interest.

### Summary of Updates

In this revised version of the manuscript, we have reorganized the Results section to improve the overall clarity and flow of the storyline. The findings are now presented in a sequence that builds more clearly from the initial observations toward the main conclusions. We have also incorporated new experimental data. These additional data help clarify points that were less well supported in the earlier version.

