## supplementary figure for "Chikungunya virus non-structural protein 2 (nsP2) inhibits RIG-I and TLR-mediated immune response"

### Slide 1
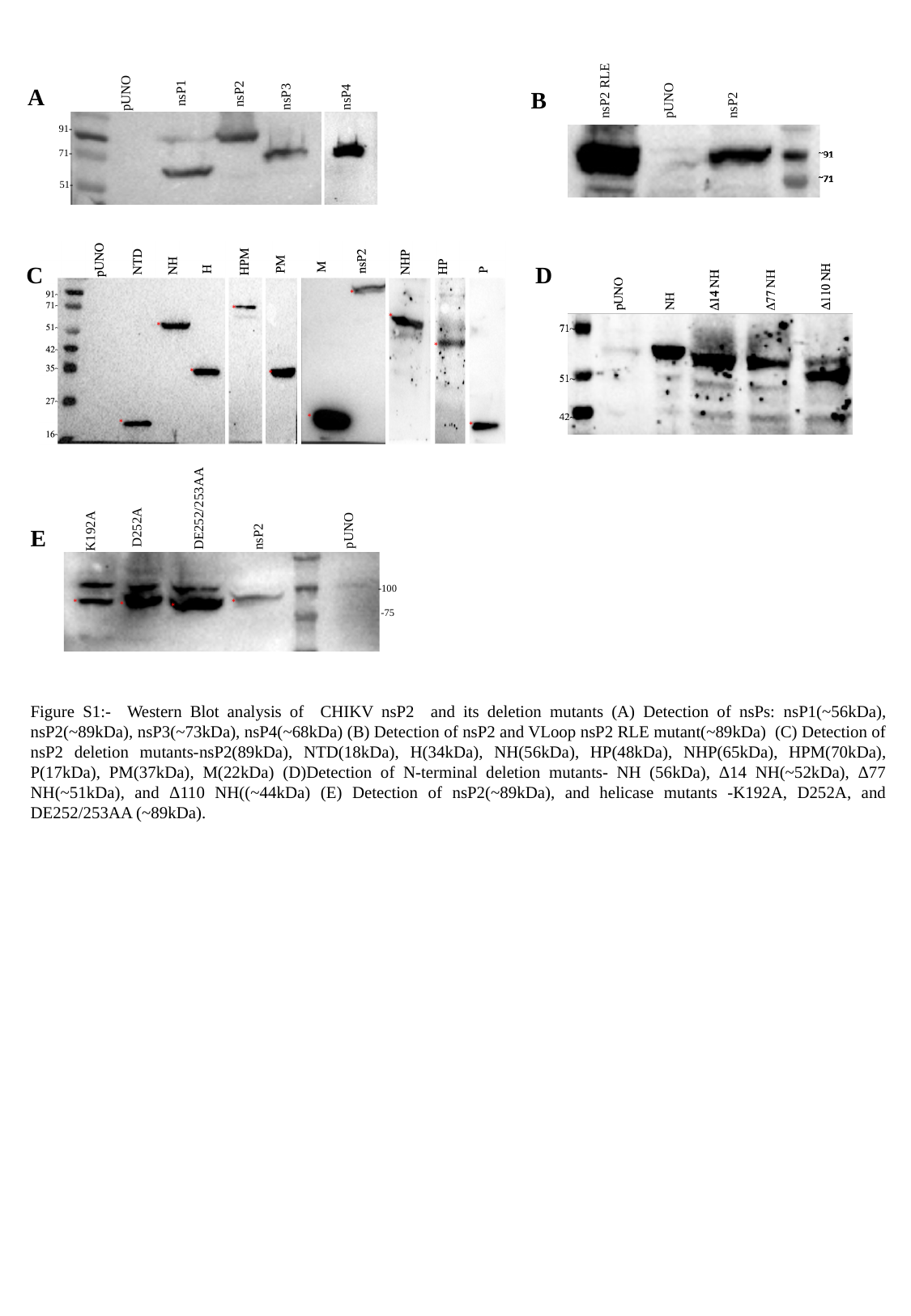

nsP2
pUNO
nsP2 RLE
nsP1
nsP2
pUNO
nsP3
91-
71-
51-
nsP4
A
B
C
D
*
*
*
*
*
*
*
*
*
*
D252A
pUNO
nsP2
DE252/253AA
K192A
-100
 -75
*
*
*
*
E
Figure S1:- Western Blot analysis of CHIKV nsP2 and its deletion mutants (A) Detection of nsPs: nsP1(~56kDa), nsP2(~89kDa), nsP3(~73kDa), nsP4(~68kDa) (B) Detection of nsP2 and VLoop nsP2 RLE mutant(~89kDa) (C) Detection of nsP2 deletion mutants-nsP2(89kDa), NTD(18kDa), H(34kDa), NH(56kDa), HP(48kDa), NHP(65kDa), HPM(70kDa), P(17kDa), PM(37kDa), M(22kDa) (D)Detection of N-terminal deletion mutants- NH (56kDa), Δ14 NH(~52kDa), Δ77 NH(~51kDa), and Δ110 NH((~44kDa) (E) Detection of nsP2(~89kDa), and helicase mutants -K192A, D252A, and DE252/253AA (~89kDa).
