## Supplementary Table for "Chikungunya virus non-structural protein 2 (nsP2) inhibits RIG-I and TLR-mediated immune response"

### Slide 1
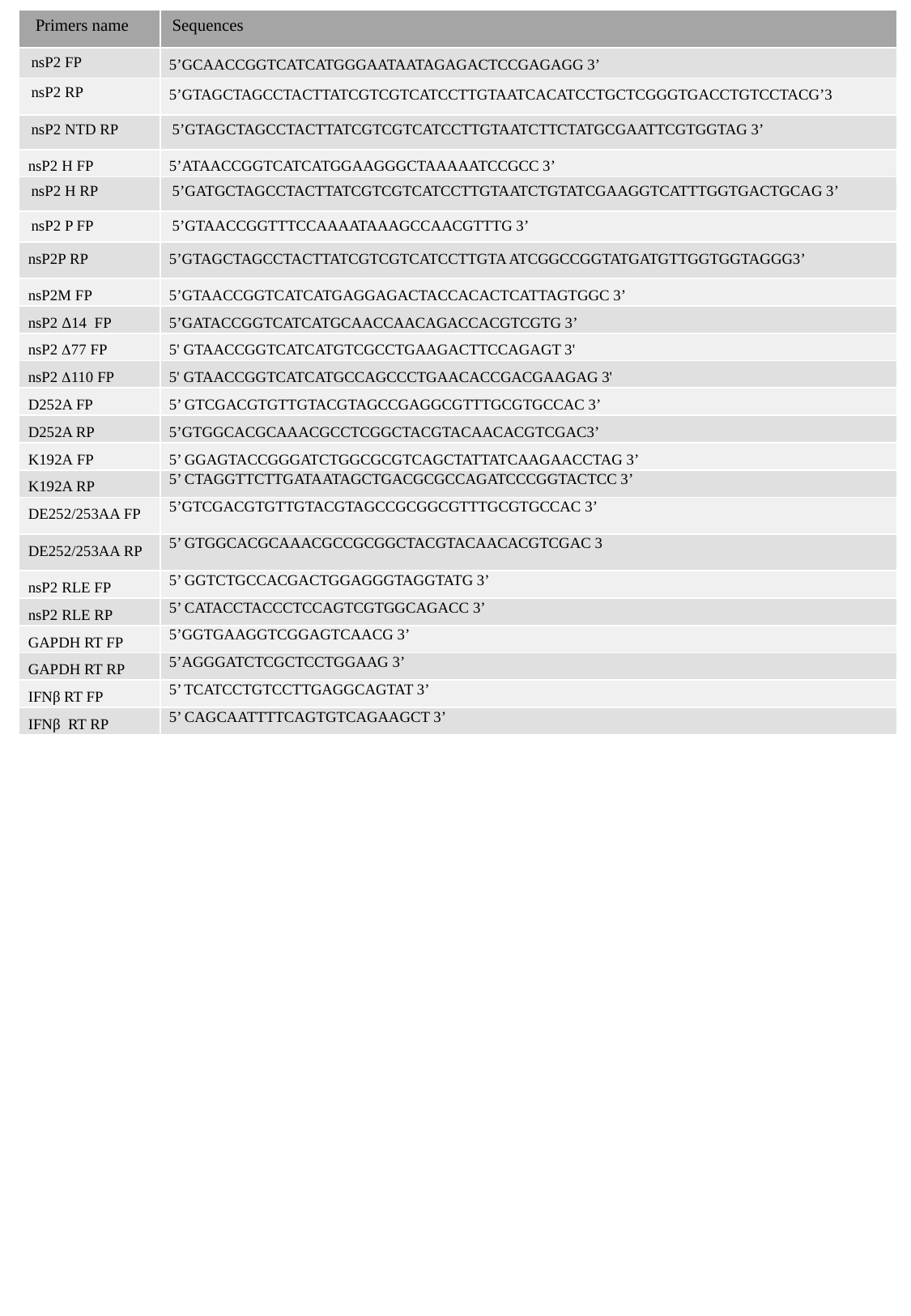

| Primers name | Sequences |
| --- | --- |
| nsP2 FP | 5’GCAACCGGTCATCATGGGAATAATAGAGACTCCGAGAGG 3’ |
| nsP2 RP | 5’GTAGCTAGCCTACTTATCGTCGTCATCCTTGTAATCACATCCTGCTCGGGTGACCTGTCCTACG’3 |
| nsP2 NTD RP | 5’GTAGCTAGCCTACTTATCGTCGTCATCCTTGTAATCTTCTATGCGAATTCGTGGTAG 3’ |
| nsP2 H FP | 5’ATAACCGGTCATCATGGAAGGGCTAAAAATCCGCC 3’ |
| nsP2 H RP | 5’GATGCTAGCCTACTTATCGTCGTCATCCTTGTAATCTGTATCGAAGGTCATTTGGTGACTGCAG 3’ |
| nsP2 P FP | 5’GTAACCGGTTTCCAAAATAAAGCCAACGTTTG 3’ |
| nsP2P RP | 5’GTAGCTAGCCTACTTATCGTCGTCATCCTTGTA ATCGGCCGGTATGATGTTGGTGGTAGGG3’ |
| nsP2M FP | 5’GTAACCGGTCATCATGAGGAGACTACCACACTCATTAGTGGC 3’ |
| nsP2 Δ14 FP | 5’GATACCGGTCATCATGCAACCAACAGACCACGTCGTG 3’ |
| nsP2 Δ77 FP | 5' GTAACCGGTCATCATGTCGCCTGAAGACTTCCAGAGT 3' |
| nsP2 Δ110 FP | 5' GTAACCGGTCATCATGCCAGCCCTGAACACCGACGAAGAG 3' |
| D252A FP | 5’ GTCGACGTGTTGTACGTAGCCGAGGCGTTTGCGTGCCAC 3’ |
| D252A RP | 5’GTGGCACGCAAACGCCTCGGCTACGTACAACACGTCGAC3’ |
| K192A FP | 5’ GGAGTACCGGGATCTGGCGCGTCAGCTATTATCAAGAACCTAG 3’ |
| K192A RP | 5’ CTAGGTTCTTGATAATAGCTGACGCGCCAGATCCCGGTACTCC 3’ |
| DE252/253AA FP | 5’GTCGACGTGTTGTACGTAGCCGCGGCGTTTGCGTGCCAC 3’ |
| DE252/253AA RP | 5’ GTGGCACGCAAACGCCGCGGCTACGTACAACACGTCGAC 3 |
| nsP2 RLE FP | 5’ GGTCTGCCACGACTGGAGGGTAGGTATG 3’ |
| nsP2 RLE RP | 5’ CATACCTACCCTCCAGTCGTGGCAGACC 3’ |
| GAPDH RT FP | 5’GGTGAAGGTCGGAGTCAACG 3’ |
| GAPDH RT RP | 5’AGGGATCTCGCTCCTGGAAG 3’ |
| IFNβ RT FP | 5’ TCATCCTGTCCTTGAGGCAGTAT 3’ |
| IFNβ RT RP | 5’ CAGCAATTTTCAGTGTCAGAAGCT 3’ |
Table S1- Primer sequences used for cloning and RT-qPCR
